# Weight loss reduces head motion: re-visiting a major confound in neuroimaging

**DOI:** 10.1101/766261

**Authors:** F. Beyer, K. Prehn, KA Wüsten, A. Villringer, J. Ordemann, A. Flöel, AV Witte

## Abstract

Head motion during magnetic resonance imaging (MRI) induces image artifacts that affect virtually every brain measure. In parallel, cross-sectional observations indicate a correlation of head motion with age, psychiatric disease status and obesity, raising the possibility of a systematic artifact-induced bias in neuroimaging outcomes in these conditions, due to the differences in head motion. Yet, a causal link between obesity and head motion has not been tested in an experimental design. Here, we show that a change in body mass index (i.e., weight loss after bariatric surgery) systematically decreases head motion during MRI. In this setting, reduced imaging artifacts due to lower head motion might result in biased estimates of neural differences induced by changes in BMI. Overall, our finding urges the need to rigorously control for within-scanner head motion to enable valid results of neuroimaging outcomes in populations that differ in head motion due to obesity or other conditions.

## Introduction

Head motion is an important confounder in neuroimaging studies of brain structure and function (Power et al., 2015;Savalia et al., 2017). Micromovements of the head, driven by spontaneous motion or respiration, strongly affect qualitative and quantitative neuroimaging outcomes, even if targeted image processing techniques are used (Parkes et al., 2018). Moreover, within-scanner head motion often correlates with the predictors under study, such as age, psychiatric disease status and obesity (Hodgson et al., 2016;Torres and Denisova, 2016;Makowski et al., 2019). This raises the possibility of a systematic image artifact-induced bias due to differences in head motion in these conditions (Satterthwaite et al., 2012;Pardoe et al., 2016).

One of the strongest predictors of head motion is body mass index (Hodgson et al., 2016;Beyer et al., 2017;Ekhtiari et al., 2019).It seems likely that physiological differences associated with higher weight, e.g., increased respiratory rate and amplitude, or spontaneous motion due to uncomfortable positioning in the magnet bore may induce this effect (Littleton, 2012). Yet, within-subject analysis indicated that differences in head motion are rather driven by a neurobiological trait which shares genetic variance with body mass index (Zeng et al., 2014;Hodgson et al., 2016). Along these lines, impulsivity, e.g. more rash action tendencies, might explain a proportion of the shared variance between head motion and obesity (Kong et al., 2014;Couvy-Duchesne et al., 2016).

Until now, mainly cross-sectional studies report on the association of BMI and head motion and little is known about how BMI changes may affect head micromovements. In this pre-registered analysis (https://osf.io/epsxt), we therefore aimed to test whether a radical weight-loss intervention (bariatric surgery) compared to a control condition induces consistent changes of head motion during MRI in obese individuals.

## Results

We compared 32 obese participants that underwent bariatric surgery to 18 obese participants in a waiting list-control condition at baseline, 6 and 12 months after surgery/waiting period (see Methods for demographics). We acquired resting-state functional MRI on a 3 Tesla Siemens MRI scanner (see Methods). Bariatric surgery compared to control condition led to a systematic reduction of within-scanner head motion, measured using framewise displacement (FD) of 150 individual brain volumes acquired with a 6 minute scan (Fig.1, linear mixed model comparison: likelihood-ratio test X^2^=11.7, df=2, p=0.0028, based on 107 observations from 50 participants). Average head motion of control participants did not change. Moreover, the magnitude of weight loss, measured as change in BMI, predicted the decrease in head motion (linear mixed model comparison, likelihood-ratio test X^2^=20.6, df=1, p<0.001, based on 107 observations from 50 participants, R^2^ of fixed effects: 0.053).

**Fig 1:**
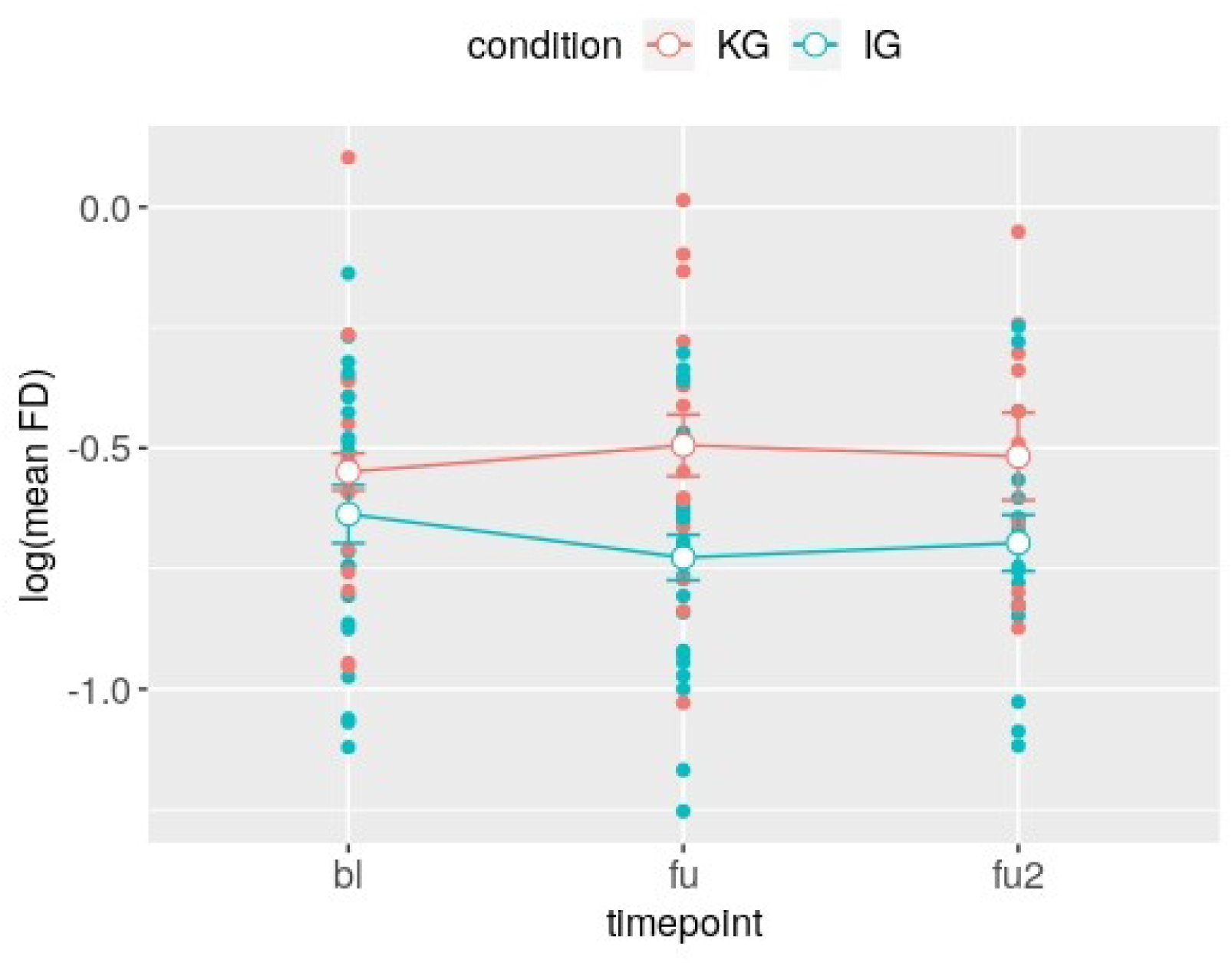
Head motion, measured as log-normalized mean framewise displacement (mean FD),decreases across the three timepoints in the intervention (IG) compared to the control condition (KG), shown in blue/red, respectively. Error bars represent within-subject errors. bl: baseline, fu: 6month follow-up, fu2: 12month follow-up.

To estimate the impact of head motion changes on structural imaging outcomes, we assume an average total grey matter volume of ∼ 600 cm^3^ and 24 cm^3^ grey matter loss per 0.1mm of mean FD increase based on results from Alexander-Bloch et al. (Alexander-Bloch et al.,2016). Thus, an estimated decrease in mean FD from baseline to follow up in the intervention group of 0.12 mm would translate into a false increase in total grey matter volume of 28.8 cm^3^ (or about 5%) after bariatric surgery, due to the decrease in head motion alone.

## Discussion

We here show that a radical change in physiological parameters, in this case body weight loss induced by bariatric surgery, reduces head micromovements during MRI. The magnitude of changes in BMI further predicted the magnitude of reduction in head motion. This indicates that head motion strongly depends on body physiology and that BMI differences may result in biased estimates of brain structure and function. Our result highlights the need of rigorous attempts to adjust for and reduce head motion during neuroimaging studies in obesity, as well as in other conditions that may be systematically related to head motion.

Our findings are in line with previous observations in several large imaging cohorts indicating that BMI accounts for 8-40% of the variance in head motion (Hodgson et al., 2016;Beyer et al.,2017) and is one of the most important predictors of head motion (Ekhtiari et al.,2019). While our estimate of a weight loss-induced bias in neuroimaging outcomes (i.e.,5% increase in total grey matter volume after bariatric surgery) relied on between-subject estimates of the effect of head motion (Alexander-Bloch et al.,2016) and therefore has to be interpreted with caution, its severity further underscores the importance to control for head motion differences in future MRI analyses. Different techniques, such as multi-echo sequences (Power et al.,2018), fixation through head molds (Power et al., 2019b) and tactile feedback during scanning (Krause et al.,2019) have been proposed to considerably reduce head motion *a priori –* which is probably the best way to handle this important confound for practically all imaging outcomes (Reuter et al.,2015;Beyer et al., 2017;Baum et al.,2018;Zhang et al.,2018).

Little is known about the mechanisms that may underlie the causal link between body weight and head motion. Possibly, obesity-related alterations in the respiratory system lead to increased real and apparent head motion in MRI scans (Littleton,2012;Power et al.,2019a).Yet, respiration rate did not mediate the association of BMI and head motion in previous studies (Hodgson et al.,2016;Ekhtiari et al.,2019). More speculative, alterations in the brain’s dopaminergic system could represent a link between BMI and head motion. Evidence suggests that BMI-related differences in dopamine receptor availability might underlie observed differences in dopamine-related functions such as reward sensitivity (Tomasi and Volkow,2013;Horstmann et al.,2015). Given that dopaminergic signaling is crucial for motor inhibition and control (Cools and D’Esposito, 2011;Robertson et al., 2015), alterations in the dopaminergic system might also influence spontaneous head motion.

A limitation of this study is that we did not assess other predictors of head motion such as impulsivity, respiration or dopaminergic function using techniques such as positron emission tomography (PET). We therefore cannot exclude that these measures mediate the observed effect. Yet, our main conclusion that physiological changes induce changes in head motion, and its implications for future studies, remain valid regardless of the exact mechanisms.

Taken together, radical weight loss induced by bariatric surgery reduces head motion in a cohort of obese individuals, indicating that the physiological state strongly determines higher head motion in obesity. This urges adequate a-priori control of head motion in neuroimaging studies of obesity and other conditions with systematically increased head motion to eliminate its confounding effects on measures of brain structure and function.

## Methods

To test whether a decrease in BMI would reduce head micro-movements in MRI, we compared 32 obese participants that underwent bariatric surgery (scanned at baseline n=22, 7m/25f, aged 41.8 ± 11.6 years, BMI: 44.2 ± 4.3kg/m^2^m (mean±SD)) to 18 obese participants in a waiting list-control (scanned at baseline n=15,7m/11 f, aged: 48.9 ± 12.1 years, BMI 42.8 ± 4.7 kg/m^2^). Resting state functional MRI (*T*^2^*-weighted EPI sequence, 150 volumes, 34 slices, repetition time 2000 ms, echo time=30 ms, flip angle=90°,voxel size=3.0 × 3.0 × 4.0mm) was acquired at three timepoints maximum (baseline, 6 and 12 months after surgery or waiting period) on a 3 Tesla Siemens Trio MRI. 11/10 participants from intervention/control group completed all three assessments and all participants were included into the linear mixed model analysis. Mean and maximal framewise displacement (FD) were calculated according to (Power et al., 2012). We deviated from the preregistration (https://osf.io/epsxt) by log-transforming mean FD values prior to the analysis. We set up a linear-mixed model with group and timepoint as fixed effect and subject as random effect using lme4 in R version 3.6.1.To test the effect of the intervention on head motion, we compared a model including the interaction of group and timepoint to a model including only main effects. We report X^2^ and p-values of the likelihood-ratio test comparing the two models. In the second analysis, we calculated within-and between-subject BMI variability and compared a full model to predict head motion to a reduced model which included only between-subject BMI. All code for this analysis is publicly available under https://github.com/fBeyer89/ADI_preproc/tree/master/head_motion_project.

## Acknowledgments

We thank T. Profitlich, S. Heßler, I.Rangus, C.Reschke, L.Kaiser, A.Winkler, L.Kerti for help in data acquisition. This work was supported by the German Research Foundation, Contract grant numbers WI 3342/3-1, Fl 379-10/1, Fl 379-11/1, FL 379-16/1, DFG-Exc257, SFB1315TPB03 and CRC1052 “Obesity mechanisms” ProjectA1.

